# The Tritryps comparative repeatome: insights on repetitive element evolution in Trypanosomatid pathogens

**DOI:** 10.1101/387217

**Authors:** Sebastián Pita, Florencia Díaz-Viraqué, Gregorio Iraola, Carlos Robello

## Abstract

The major human pathogens *Trypanosoma cruzi*, *Trypanosoma brucei* and *Leishmania major* are collectively known as the Tritryps. The initial comparative analysis of their genomes has uncovered that Tritryps share a great number of genes, but repetitive DNA seems to be extremely variable between them. However, the in-depth characterization of repetitive DNA in these pathogens has been in part neglected, mainly due to the well-known technical challenges of studying repetitive sequences from *de novo* assemblies using short reads. Here, we compared the repetitive DNA repertories between the Tritryps genomes using genome-wide, low-coverage Illumina sequencing coupled to RepeatExplorer analysis. Our work demonstrates that this extensively implemented approach for studying higher eukaryote repeatomes is also useful for protozoan parasites like trypanosomatids, as we recovered previously observed differences in the presence and amount of repetitive DNA families. Additionally, our estimations of repetitive DNA abundance were comparable to those obtained from enhanced-quality assemblies using longer reads. Importantly, our methodology allowed us to describe a previously unknown transposable element in *Leishmania major* (TATE family), highlighting its potential to accurately recover distinctive features from poorly characterized repeatomes. Together, our results support the application of this low-cost, low-coverage sequencing approach for the extensive characterization of repetitive DNA evolutionary dynamics in trypanosomatid and other protozooan genomes.

## Main Text

Collectively known as the “Tritryps”, the unicellular mono-flagellated protozoan parasites *Trypanosoma cruzi, Leishmania major* and *Trypanosoma brucei* are the causative agents of American trypanosomiasis, cutaneous leishmaniasis and African trypanosomiasis, respectively. These dixenous parasites belong to the family *Trypanosomatidae*, within the order *Kinetoplastida* (Votýpka et al. 2015). Despite Tritryps share many general characteristics which are used as distinctive taxonomic markers (*i.e.* their unique mitochondria known as kinetoplast), each species has its own insect vector, particular life-cycle features, different target tissues, and distinct disease pathogenesis in mammalian hosts (El-Sayed et al. 2005a; Jackson 2015).

The genomes of *T. cruzi*, *L. major* and *T. brucei* have been initially sequenced and compared to better understand gene evolution and genetic variation in these related pathogens (El-Sayed et al. 2005a; Ghedin et al. 2004). A remarkable finding derived from the comparative analysis of Tritryps genomes was the great number of shared genes (El-Sayed et al. 2005a). However, the repetitive DNA was extremely different in these species. Repetitive DNA sequences are scarce in the *L. major* genome, but comprises up to half of the *T. cruzi* genome. Moreover, *L. major* is believed to be devoid of active transposable elements (TEs) (Ghedin et al. 2004; Ivens et al. 2005; Bringaud et al. 2006), but both *T. cruzi* and *T. brucei* genomes harbor intact and autonomous TEs (Wickstead et al. 2003; El-Sayed et al. 2005b; Bringaud et al. 2008; Thomas et al. 2010; Berná et al. 2018).

Genome annotation procedures are mainly focused on standard genetic elements, frequently neglecting repetitive sequences due to their hard-achieving *de novo* assembly (Treangen & Salzberg 2011). As a consequence, repetitive DNA is poorly described and studied (Altemose et al. 2014). In this context, RepeatExplorer has emerged as a widely used approach to comprehensively evaluate the nature of repetitive sequences. This bioinformatic tool attempts to cluster genome-wide, low-coverage sequencing reads using a graph-based algorithm to characterize and quantify the complete repetitive DNA fraction of a genome (Novák et al. 2010, 2013, 2017), which nowadays is known as the “repeatome” (Maumus & Quesneville 2014). Low-coverage sequencing is a cost-effective approach that does not require having previous information about the target genome and avoids dealing with whole-genome assemblies. Beyond RepeatExplorer was originally conceived to analyze plant repeatomes, it has been successfully applied in mammals (Pagán et al. 2012), insects (Ruiz-Ruano et al. 2016; Palacios-Gimenez et al. 2017; Pita et al. 2017) and fishes (Utsunomia et al. 2017). Here, we used low-coverage sequencing and the RepeatExplorer approach to compare the repeatomes of *T. cruzi*, *L. major* and *T. brucei* seeking for insights into the evolution of repetitive sequences in these pathogenic parasites.

First, kinetoplast DNA (mini and maxicircles) was removed from raw Illumina reads and after quality filtering a random subsampling was performed to obtain ~1x coverage in each genome. This resulted in 353,334 reads from *T. cruzi*, 173,836 reads from *T. brucei* and 218,778 reads from *L. major* that were subsequently used in the RepeatExplorer analyses. The software initially identified 293, 203 and 199 clusters for *T. cruzi*, *T. brucei* and *L.major*, respectively. In *T. cruzi* we estimated that 51.25% of the genome corresponds to repetitive DNA sequences. Out of them, 28.81% were annotated as coding sequences belonging to multigenic families, 8.85% as LINEs (Long Interspersed Elements), 3.73% as DIRS-like or tyrosin recombinase (YR) elements (mostly VIPER element), 3.48% as satellite DNA, 0.31% represented ribosomal DNA (rDNA) and 5.07% remained as undetermined repeats. Conversely, in *T. brucei* only 20.69% of the genome harbors repetitive DNA sequences. Out of them, we were able to determine that 9.53% belong to coding sequences from multigenic families, 5.67% to LINE TEs, 3.59% were satellite DNA repeats, 0.33% as rDNA and 1.57% of the genome remained as undetermined repeats. Finally, the repetitive DNA fraction in *L. major* was smaller than in the genus *Trypanosoma*, corresponding only to 8.80% of the genome. The vast majority of this repetitive DNA consisted in multigenic families (see details below), which reached the 3.93% of the genome. Additionally, 1.32% was identified as TEs named telomere-associated mobile elements (TATEs), 0.29% as LINE TEs and 0.27% assigned to satellite DNA repeats. Also, several clusters belonged to rDNA genes and snoRNA regions, which accounted for the 0.57% and 0.34% of the genome, respectively. The remaining 2.09% of the genome was annotated as undetermined repeats (Fig. 1).

**Figure 1.**
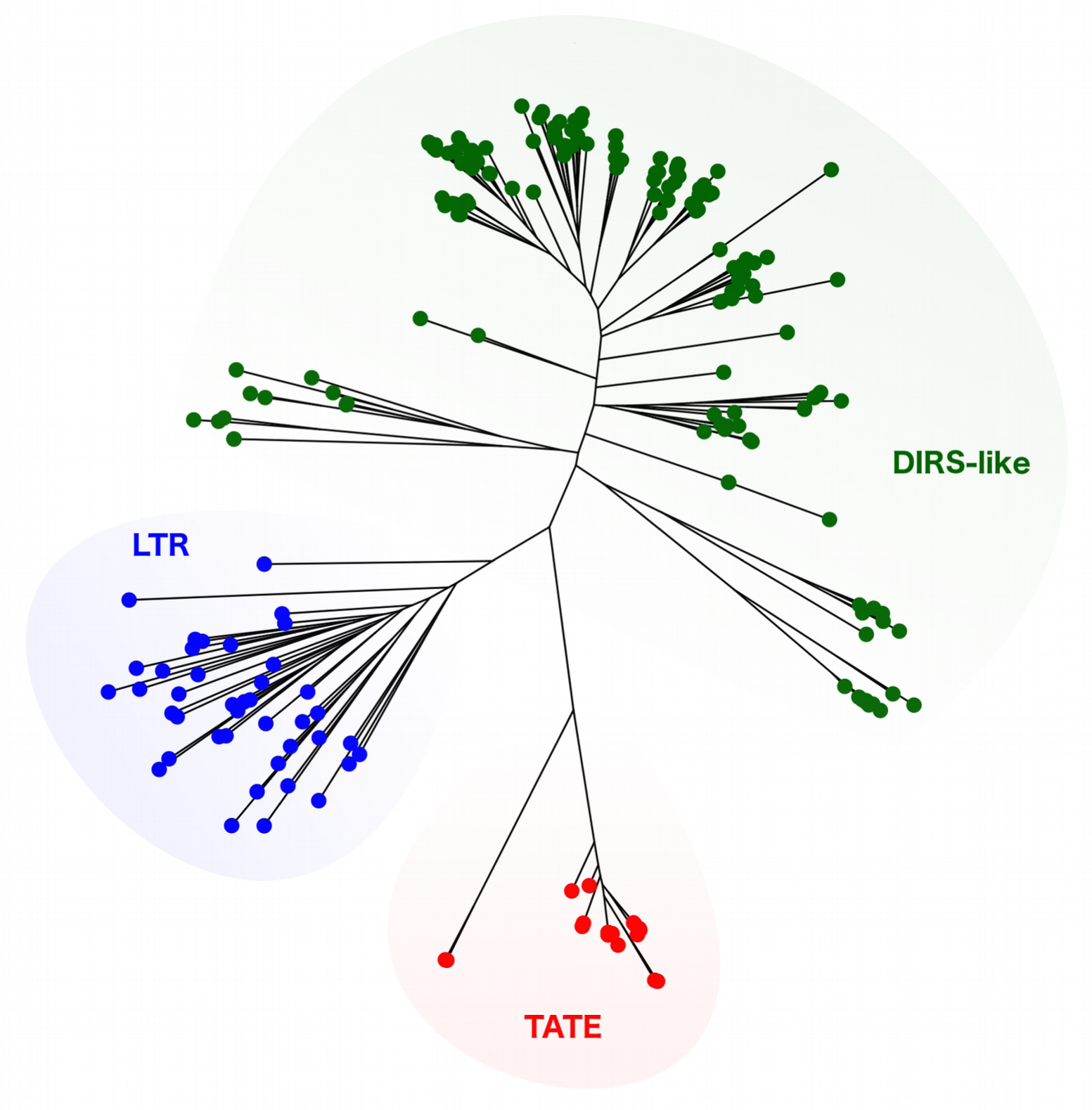
Comparison of *T. cruzi*, *T. brucei* and *L. major* repeatomes. Bar plots show the relative amount of each repetitive DNA fraction on the (A) *T. cruzi*, (B) *T. brucei* and (C) *L. major* genome. Pie charts represent the relative amount of repetitive and non-repetitive DNA on each genome.

In agreement to previous quantifications of repetitive DNA in the genomes of *T. cruzi* CL Brener (El-Sayed et al. 2005b) and X-10 Sylvio (Franzén et al. 2011) strains our current analysis using *T. cruzi* Dm28c showed that almost half of the genome is composed by these sequences. In addition, our estimations of repetitive DNA are consistent with the new reference genomes of *T. cruzi* based on long-read sequencing, which used TCC and Dm28c as well (Berná et al. 2018). Within the repetitive fraction, the most abundant sequences correspond to multigenic families as previously described for the CLBrener and X-10 Sylvio strains, being this a salient feature in the *T. cruzi* genome. However, the relative abundance of each family remained uncertain and probably underestimated (El-Sayed et al. 2005b). Again, here we accurately compared our estimations with the new non-collapsed *T. cruzi* reference genomes (Berná et al. 2018), obtaining quite similar quantifications of the multigenic families as a whole. Additionally, we were able to individually quantify the relative abundance of trans-sialidase (TS), retrotransposon hot spot (RHS), mucins, mucin-associated surface proteins (MASP), surface protein dispersed gene family-1 (DGF-1) and GP63 multigenic families (Fig. 1). The amount of these families vary among different *T. cruzi* strains, as has been shown between CLBrener and X-10 Sylvio, which could be related to their infection capacity, since these protein-coding genes are involved in the parasite-host interactions (Franzén et al. 2011). The application of low-coverage sequencing and RepeatExplorer analysis over multiple *T. cruzi* strains with differential infectivity may uncover the relationship between multigenic dynamics and pathogenesis. TEs ranked second in terms of abundance being almost 13% of the genome. LINEs (such as the L1Tc, NARTc, CZAR, and TcTREZO elements) where more abundant than YR elements (VIPER element and their non-autonomous derivative SIRE). These repetitive DNA sequences were also underestimated on previous analyses (El-Sayed et al. 2005b; Franzén et al. 2011; Berná et al. 2018). TEs richness differences between strains has been already described (Vargas et al. 2004) and could be attributed to natural variations between *T. cruzi* strains, but considering the remarkable disparity (5% estimated for CL Brener) this must deserve further attention, since additional factors than strain diversification may be explaining TEs dynamics. Moreover, TEs quantification on both newly generated reference genomes showed almost the same TEs abundance for TCC strain -closely related to CL Brenner- and Dm28c strain (Berná et al. 2018), evidencing that RepeatExplorer is a valid tool for TEs recognition and quantification. Lastly, satellite DNA sequences encompass more than 3% of the genome, being vastly dominated by the 195nt satellite. Previous 195nt satellite quantification on CL Brenner estimated that 5% of the genome is composed by this repeat (Martins et al. 2008). However, variation of 195nt abundance has been reported to be four to six fold between DTU TcI and DTU TcII strains (Elias et al. 2003; Vargas et al. 2004). This difference is also observed in the new reference genomes from TCC and Dm28c. Actually, the Dm28c quantification is close to that reported here, reinforcing that low-coverage sequencing provides reproducible estimations of repeat element abundances. Several other tandem repeats have been recently described (Berná et al. 2018) but only a few of them were retrieved by RepeatExplorer, indicating that their abundances are below the threshold set for a standard analysis. However, we aimed to render a coarse-grain, genome-wide overview rather than a meticulous description of all repeats.

*T. brucei* genome is composed by around 20% of repetitive DNA. Similar to *T. cruzi*, multigenic families were the most abundant repeats reaching around 10% of the genome, with RHS and VGS/ESAG as the most representative families. TEs in *T. brucei* represented 5.67% of the genome, but in this case is only composed by LINE sequences, such as the *Tbingi* elements, its related non-autonomous RIMEs, and a few SLAC elements. Although VIPER elements are described in *T. brucei* (Lorenzi et al. 2006), these repeats are known to be in very low copy number, hence undetectable under our approach. The first draft genome of *T. brucei* presented in 2005 (Berriman et al. 2005) only reported that subtelomeric genes were just over 20% and that TEs represented 2% of the genome (El-Sayed et al. 2005b), however nothing is said about the satellite DNA. Our results showed two prominent satellite DNA families, the 177-bp repeat described to be part of intermediate and minicromosomes which are enriched by VSG genes (Sloof et al. 1983; Wickstead et al. 2004; Obado et al. 2005), and the 147 bp repeat (named CIR147) present in the centromere form the majority of macrochromosomes (Obado et al. 2007).

The most surprising results came along with the TEs analysis in *L. major*. Repetitive DNA comprehends less than 10% of the genome, as was expected since former genome analyses described smaller subtelomeric regions than in *Trypanosoma* species (Ivens et al. 2005; Peacock et al. 2007). Furthermore, the closely related *L. braziliensis* and *L. infantum* have also around 10% of the genome composed by DNA repeats (Peacock et al. 2007). Although the reference genome for *L. infanum* has been re-sequenced using long reads technology, revealing an expansion of coding genes copy number, the amount of repetitive DNA was not cited (González-De La Fuente et al. 2017). As observed in *Trypanosoma*, the majority of the repeated genome was represented by gene-coding sequences, being GP-63 and the Leucin-rich-repeats among the most abundant elements. Remarkably, as in *L. braziliensis* genome (Peacock et al. 2007) but not described so far for *L. major*, we found traces of a the LINE element related to CRE2 (from *Crithidia fasciculata*), which is also related with CZAR and SLACs TEs from *T. cruzi* and *T. brucei*, respectively. Another interesting finding was the presence of a truncated element bearing a reverse transcriptase domain, from the LINE order. This probably corresponds to the LmDIRE elements, which are included in the *ingi2* clade (Bringaud et al. 2009). By far, an exceptional finding was the recovering of TATE copies representing 1.32% of the genome. These elements were previously reported in other *Leishmania* species from the subgenus *Viannia*, such as *L. braziliensis* (Peacock et al. 2007) and *L. panamensis* (Llanes et al. 2015), but not from the subgenus *Leishmania*, as *L. major*. Sequence similarity searches on the *L. major* genome available on TriTrypdb (https://www.TriTrypdb.org) did not retrieve any positive results. Currently, TATEs are not classified within any of the TEs families, nor even as a concrete class. However, the presence of a tyrosine recombinase suggests that possibly TATEs are DIRS-like TEs (Peacock et al. 2007). Here we were able to reconstruct a consensus sequence for the retrotranscriptase domain of the *L. major* TATE element, and determine that all kinetoplastid TATEs described hitherto form a separated clade from other DIRS-like elements (Fig. 2). Further analysis on these elements would be of major interest for better understanding the dynamics of *Leishmania* genomes. Beyond the already know impact of TEs in trypanosomatid genomes (Bringaud et al. 2008; Thomas et al. 2010), our finding that TATE elements account for a considerable part of the *L. major* genome, could change the evolutionary paradigm of a genome that was believed to be almost TE-free. Actually, it has been already suggested that TATEs are not restricted to telomeric regions in *L. panamensis* genome, and that they could be playing a central role in gene regulation and structuring (Llanes et al. 2015). For example, being candidates to participate on recombinational events leading to genetic amplification (Ubeda et al. 2014).

**Figure 2.**
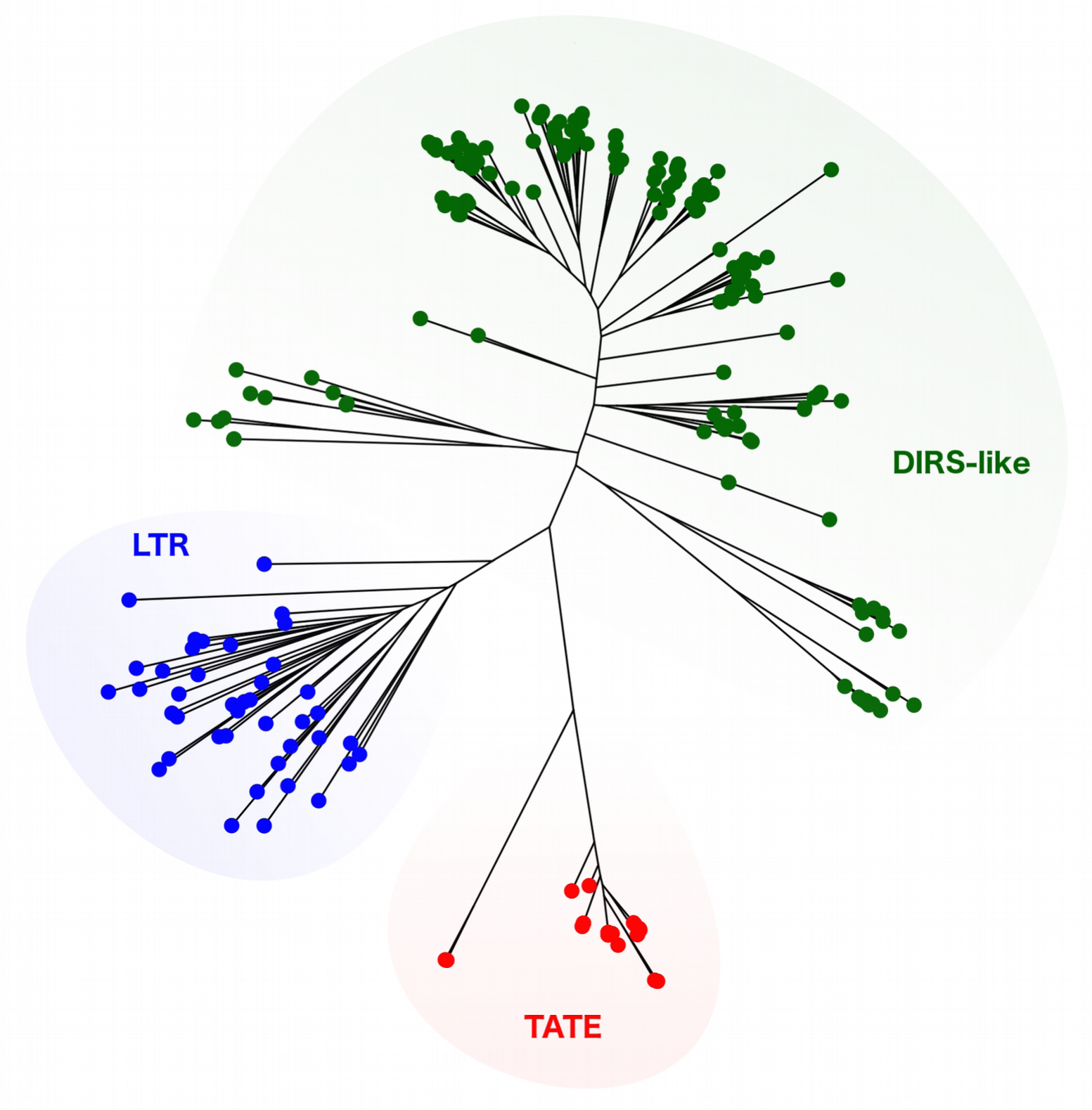
Phylogenetic characterization of TATE elements. Maximum-Likelihood phylogenetic tree using full retrotranscriptase domain sequences from TATE and other related elements (LTR and DIRS-like) recovered from a wide taxonomic representation of eukaryote organisms.

In conclusion, we have shown that our results are comparable to those obtained for other Tritryps strains and implementing different sequencing strategies, such as high-coverage and long-read genomic assemblies. This supports that our method using low-coverage, Illumina short reads is useful for a genome-wide characterization of trypanosomatid repeatomes, and could be useful to perform comparative analyses of the repetitive DNA repertories in other protozoan species. Noteworthy, our strategy allowed to identify genetic features that were not described so far, such as TATEs elements in the *L. major* genome.

## Materials and Methods

### Strains and DNA purification

*T. cruzi* Dm28c (Contreras et al. 1988) epimastigotes were cultured axenically in liver infusion tryptose medium supplemented with 10% (v/v) inactivated fetal bovine serum (*GIBCO*) at 28°C. *L major* (Instituto Oswaldo Cruz – Fiocruz, IOCL 2821) and *T. brucei* were cultured in modified RPMI medium containing 10% (v/v) inactivated fetal bovine serum (*GIBCO*) at 28°C. Quick-DNA Universal kit (Zymo Research, Irvine, California, USA) was used according to the manufacturer’s specifications for isolation of genomic DNA in logarithmic growth phase. The DNA was resuspended in sterile distilled water and stored at 4 °C until use. Quantification was performed using Qubit™ dsDNA HS Assay Kit (Invitrogen by Thermo Fisher Scinetific, USA).

### Illumina sequencing and bioinformatic analyses

Genomic libraries were prepared with the Nextera XT DNA Sample Preparation Kit (*Illumina*), analyzed using 2100 Bioanalyzer (*Agilent*) and then sequenced using a MiSeq Illumina platform, which produced 540831, 1790895 and 1322286 pair-end reads (2×150 cycles) for *T. cruzi*, *T. brucei* and *L.major*, respectively. Graph-based clustering analyses were carried on separately using RepeatExplorer, implemented within the Galaxy environment (http:// repeatexplorer.org/) (Novák et al. 2010, 2013, 2017).

*L. major* TATE consensus sequence was determined assembling the raw reads wich belonged to RepeatExplorer clusters annotated as TATEs. Assembly was performed using CAP3 (Huang & Madan 1999) and sequence alignment using SeaView (Gouy et al. 2010). DIRS-like sequences were downloaded from Repbase (https://www.girinst.org/repbase/) and only those with complete retrotranscriptase domains were used. DIRS-1 retrotrascriptase domain sequences were also recovered from the GenBank cd03714 sequence cluster, and retrotrascriptase domain sequences from other LTR elements were used as outgroup (cd01647 sequence cluster). Alignment of amino acid sequences was performed using MAFFT software (Katoh et al. 2002) under the G-INS-i method. Phylogenetic reconstruction was performed with PhyML (Guindon et al. 2010) under the WAG substitution model and the aLRT (Shimodaria-Hasegawa-like) test was employed for internal node support.

## Acknowledgments

This work was funded by Agencia Nacional de Investigación e Innovación (ANII) DCI-ALA/2011/023–502, ‘Contrato de apoyo a las políticas de innovación y cohesión territorial’, and ‘Fondo para la Convergencia Estructural del Mercado Común del Sur (FOCEM)’ 03/11. SP, GI and CR are members of the ‘Sistema Nacional de Investigadores (ANII)’; FDV has a ANII doctoral fellowship No. POS_NAC_2016_1_129916.

